# Comparative genome analysis of a multidrug-resistant *Pseudomonas aeruginosa* sequence type 277 clone that harbours two copies of the *bla*_SPM-1_ gene and multiple single nucleotide polymorphisms in other resistance-associated genes

**DOI:** 10.1101/693440

**Authors:** Ana Paula Barbosa do Nascimento, Fernando Medeiros Filho, Hério Sousa, Hermes Senger, Rodolpho Mattos Albano, Marcelo Trindade dos Santos, Ana Paula D’Alincourt Carvalho-Assef, Fabrício Alves Barbosa da Silva

**Affiliations:** Programa de Computação Científica, Fundação Oswaldo Cruz, Rio de Janeiro, Rio de Janeiro, Brazil; Departamento de Computação, Universidade Federal de São Carlos, São Carlos, São Paulo, Brazil; Departamento de Bioquímica, Universidade do Estado do Rio de Janeiro, Rio de Janeiro, Rio de Janeiro, Brazil; Laboratório Nacional de Computação Científica, Petrópolis, Rio de Janeiro, Brazil; Laboratório de Pesquisa em Infecção Hospitalar, Fundação Oswaldo Cruz, Rio de Janeiro, Rio de Janeiro, Brazil

## Abstract

*Pseudomonas aeruginosa* is one of the most common pathogens related to healthcare-associated infections. The Brazilian isolate, named CCBH4851, is a multidrug-resistant clone belonging to the sequence type 277. The antimicrobial resistance mechanisms of the CCBH4851 strain are associated with the presence of *bla*_SPM-1_ gene, encoding a metallo-beta-lactamase, in addition to other exogenously acquired genes. Whole-genome sequencing studies focusing on emerging pathogens are essential to identify physiological key aspects that may lead to the exposure of new targets for therapy. This study was designed to characterize the genome of *Pseudomonas aeruginosa* CCBH4851 through the detection of genomic features and genome comparison with other *Pseudomonas aeruginosa* strains. The CCBH4851 closed genome showed features that were consistent with data reported for the specie. However, comparative genomics revealed the absence of genes important for pathogenesis. On the other hand, CCBH4851 genome contained acquired genomic islands that carry additional virulence and antimicrobial resistance-related genes. The presence of single nucleotide polymorphisms in the core genome, mainly those located in resistance-associated genes, suggests that these mutations could influence the multidrug-resistant behavior of CCBH4851. Overall, the characterization of *Pseudomonas aeruginosa* CCBH4851 complete genome revealed several features that could directly impact the profile of virulence and antibiotic resistance of this pathogen in infectious outbreaks.

## Introduction

*Pseudomonas aeruginosa* is one of the most common pathogens related to healthcare-associated infections in hospitalized individuals worldwide. Multidrug-resistant (MDR) isolates, particularly those non-susceptible to carbapenems, have become the major concern of health institutions. Overall, the carbapenem resistance is given by genes encoding a rising class of beta-lactamases, the metallo-beta-lactamases (MBLs), associated with other intrinsic resistance mechanisms. In addition, *P. aeruginosa* has a remarkable ability to acquire exogenous genes conferring resistance to carbapenems and other antibiotic classes [1]. The capacity of *P. aeruginosa* to grow in biofilms is an aggravating factor which can play an important role in protecting the bacteria from chemotherapy [2]. In Brazil, a recent report shows 42.9% of the *P. aeruginosa* isolates recovered from primary bloodstream infections among adult patients hospitalized in intensive care units were resistant to carbapenems [3]. The main carbapenem resistance mechanism associated with Brazilian isolates is the production of a MBL denominated SPM-1, which confers broad-spectrum resistance to all beta-lactams except for aztreonam. In 2002, the SPM-1-encoding gene (*bla*_SPM-1_) was first identified in a clinical isolate recovered from a blood culture of a patient admitted in a hospital located in the state of São Paulo. Over the past years, several isolates carrying *bla*_SPM-1_ were recovered from multiple *P. aeruginosa* hospital infection outbreaks widespread in the Brazilian territory. Multilocus sequence typing included most of these isolates within the sequence type 277 (ST-277) which contains *P. aeruginosa* strains from different countries. However, *bla*_SPM-1_ presence was only detected in isolates originating from Brazil and it is unknown why the SPM-1 epidemiology is restricted to a specific region while other MBLs tend to spread worldwide. Nevertheless, the risk of SPM-1 worldwide dissemination should not be neglected [4]. In 2008, an isolate was recovered from the catheter tip of a patient admitted in a hospital located in the state of Goiás. The strain, named CCBH4851, was submitted to antimicrobial susceptibility assays. Among the agents tested, this strain was resistant to aztreonam, amikacin, gentamicin, ceftazidime, cefepime, ciprofloxacin, imipenem, meropenem, piperacillin-tazobactam, being susceptible only to polymyxin B [5]. Multidrug resistance scenarios like those presented by CCBH4851 highlight the urgent need for the development of new antibiotics. Thereby, recent whole-genome sequencing studies are focusing on MDR *P. aeruginosa* emerging pathogens in order to identify key aspects of their physiology that may lead to the exposure of new targets for therapy. This study was designed to thoroughly characterize the complete closed genome of *P. aeruginosa* CCBH4851 through the detection of genomic features and genome comparison with other *P. aeruginosa* strains. This study intends to be a reference for future research using CCBH4851 as a model organism.

## Materials and methods

### Bacterial strains

The focus of the present study was the bacterial strain *Pseudomonas aeruginosa* CCBH4851. The isolate is available in the Coleção de Culturas de Bactérias de Origem Hospitalar (CCBH) located at Fundação Oswaldo Cruz (WDCM947; CGEN022/2010). Other strains used to perform comparative genome analysis are listed in Table 1.

**Table 1.** Bacterial strains used for comparative genome analysis.

| Strain | No. of contigs | Total size | Sequence type | Accession no. |
| --- | --- | --- | --- | --- |
| *- Group A* |  |  |  |  |
| PAO1 | 1 | 6,264,404 | 549 | NC_002516 |
| UCBPP-PA14 (PA14) | 1 | 6,537,648 | 253 | NC_008463 |
| *- Group B* |  |  |  |  |
| PA1088 | 1 | 6,721,480 | 277 | CP015001 |
| PA3448 | 2 | 6,794,242 | 277 | LVWC01000000 |
| PA7790 | 1 | 7,018,690 | 277 | CP014999 |
| PA8281 | 1 | 6,928,736 | 277 | CP015002 |
| PA11803 | 1 | 7,006,578 | 277 | CP015003 |
| PA12117 | 3 | 6,643,782 | 277 | LVXB01000000 |
| *- Group C* |  |  |  |  |
| PA7 | 1 | 6,588,339 | 1195 | NC_009656 |
| NCGM2.S1 | 1 | 6,764,661 | 235 | AP012280 |
| WH-SGI-V-07170 | 119 | 6,813,449 | 235 | NZ_LLLZ00000000 |
| WH-SGI-V-07694 | 106 | 6,776,438 | 235 | NZ_LLSO00000000 |
| AZPAE15021 | 129 | 6,686,424 | 244 | NZ_JTNZ00000000 |
| WH-SGI-V-07409 | 144 | 6,932,716 | 244 | NZ_LLOW00000000 |
| WH-SGI-V-07705 | 130 | 6,802,755 | 244 | NZ_LLSZ00000000 |

### Genome sequencing, reassembly and re-annotation

The whole-genome sequencing of CCBH4851 was performed using the Illumina MiSeq platform. Genome assembly resulted in a draft genome comprising 150 contigs which were annotated as described in previous work [5]. Later, an additional whole-genome sequencing was performed using the PacBio platform. *De novo* assembly was carried out using the MaSuRCA assembler which allowed the combination of Illumina short reads with PacBio long reads resulting in an improvement of the original assembly [6]. A single contig was obtained and was annotated using a customized pipeline. First, annotation from the closely related strain *Pseudomonas aeruginosa* PAO1 was transferred to CCBH4851 genome using the Rapid Annotation Transfer Tool software [7]. Then, coding sequences (CDSs) were predicted by GeneMarkS [8]. Prediction of rRNAs and tRNAs were performed by RNAmmer and tRNAscan-SE software, respectively [9, 10]. The predicted CDSs were functionally annotated based on homology searches against public databases, such as NR, COG, KEGG, UniProt and InterPro. Finally, the annotations from previous steps were compared, combined and manually curated. Additionally, PRIAM software was used to assign an enzyme commission number to CDSs [11]. All genome visualization was made using the Artemis software [12]. The circular plot of CCBH4851 genome was produced by Circos software [13]. Final complete genome annotation has been deposited in Genbank under the accession number CP021380.

### Comparative genome analysis

Whole-genome comparison was performed to identify similarities and differences between CCBH4851 and each group of strains listed in Table 1. The genomes were analyzed using the bidirectional best-hit (BBH) clustering method based on homology searches using the BLAST algorithm to identify pairs of corresponding genes that are each other best hit when different genomes are compared [14]. All-against-all BLAST alignments were performed using the following parameters: ≥90% coverage, ≥90% similarity and E value cut-off of 1e-10. An R algorithm was applied over BLAST results to identify the BBHs [15]. Core, accessory and unique genomes were analyzed using a MySQL database created with the data generated in previous steps. The CDSs were classified into Clusters of Orthologous Groups (COG) funtional categories by eggNOG-Mapper web application [16]. The identification of genomic islands was carried out by IslandViewer [17]. Insertion sequences (ISs) were identified using the ISSaga web application [18]. Presence of CRISPR-Cas system was assessed by CRISPRCasFinder web application [19]. In order to detect genes involved with antimicrobial resistance mechanisms, the protein sequences of CCBH4851 genome were used to perform BLAST searches against The Comprehensive Antibiotic Resistance Database using the Resistance Gene Identifier web application [20]. Also, the protein sequences were searched against the Virulence Factors Database of Pathogenic Bacteria [21] using BLASTP with ≥75% coverage, ≥50% identity and E value cut-off of 1e-10. Regulatory proteins were predicted using the Predicted Prokaryotic Regulatory Proteins web application [22]. Detection of single nucleotide polymorphisms (SNPs) was performed by Snippy software based on the alignment of unassembled reads with the *P. aeruginosa* PAO1 genome sequence using default parameters [23]. The effect of non-synonymous variants were analyzed using the HOPE web application [24].

## Results

### Genome features of *P. aeruginosa* CCBH4851

A complete genome sequence was obtained by the combined assembly of Illumina short and PacBio long reads. The genome consisted of a single circular chromosome with 6,834,257 bp and a G+C content of 66.07%. The number of CDSs was 6,211 with an average length of 976 bp, which represents 88.69% of the genome. COG families were attributed to 5,247 CDSs comprising 20 functional categories distributed along the chromosome (Fig 1). A summary of the genomic features found in CCBH4851 complete genome is listed in Table 2.

**Table 2.** Genome features of *P. aeruginosa* CCBH4851 compared to *P. aeruginosa* PAO1 reference strain.

| Feature | Strain |  |
| --- | --- | --- |
|  | CCBH4851 | PAO1 |
| Genome status | complete | complete |
| Source | catheter tip | wound |
| Country | Brazil | Australia |
| Year | 2008 | unknown |
| Genome size (bp) | 6,834,257 | 6,264,404 |
| G+C content (%) | 66.08 | 66.56 |
| No. of total genes | 6,319 | 5,697 |
| No. of pseudogenes | 78 | 19 |
| No. of total CDSs | 6,211 | 5,572 |
| No. of hypothetical proteins | 2,476 | 2,256 |
| No. of rRNAs | 13 | 13 |
| No. of tRNAs | 64 | 63 |
| No. of other RNAs | 29 | 30 |

**Fig 1.**
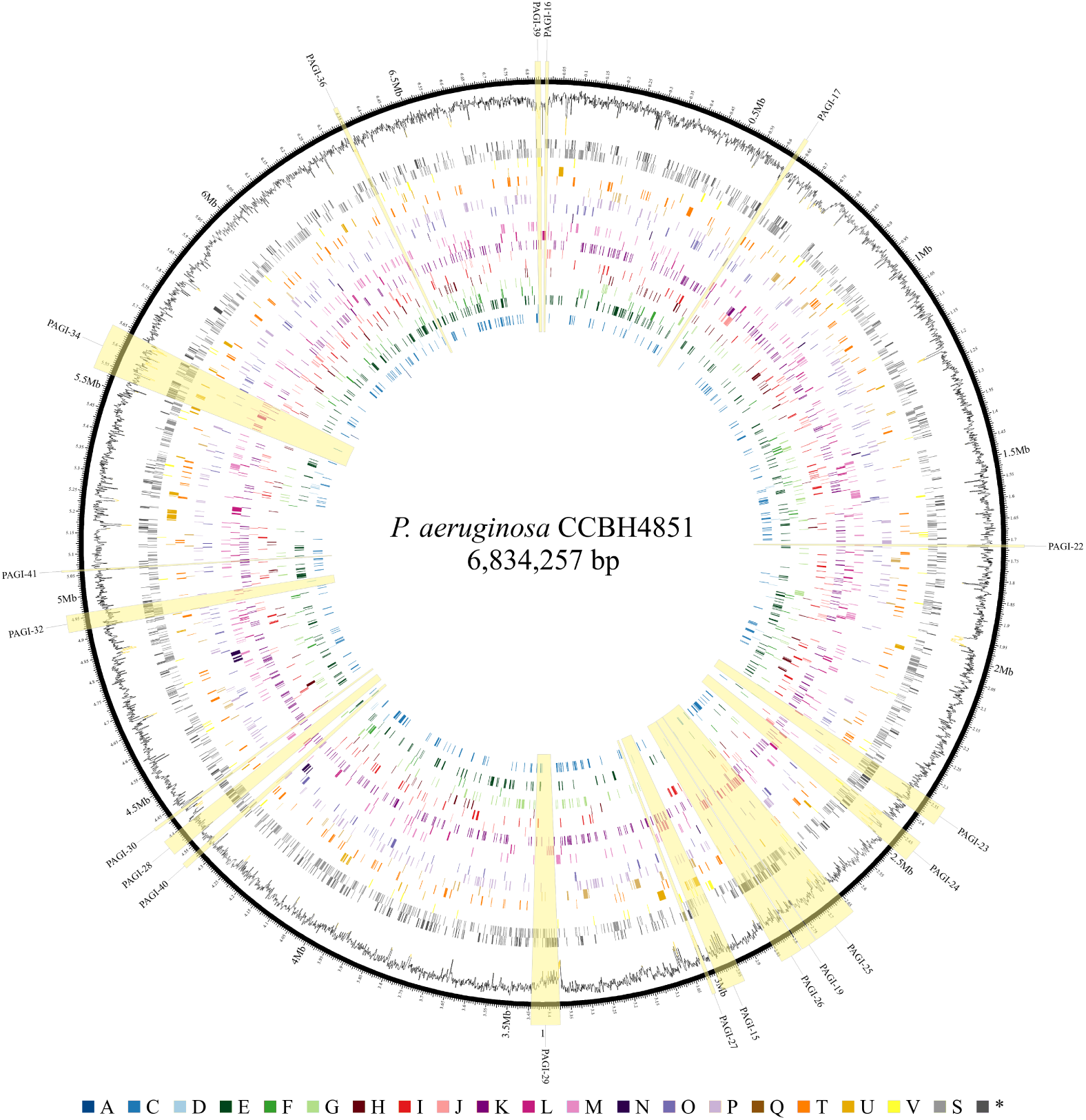
Circular representation of *P. aeruginosa* CCBH4851 genome. From the inside out, the rings display the distribution of genes along the chromosome based on COG classification: (A) RNA processing and modification; (C) energy production and conversion; (D) cell cycle control, cell division, chromosome partitioning; (E) amino acid transport and metabolism; (F) nucleotide transport and metabolism; (G) carbohydrate transport and metabolism; (H) coenzyme transport and metabolism; (I) lipid transport and metabolism; (J) translation, ribosomal structure, and biogenesis; (K) transcription; (L) replication, recombination, and repair; (M) cell wall, membrane, and envelope biogenesis; (N) cell motility; (O) post-translational modification, protein turnover, and chaperones; (P) inorganic ion transport and metabolism; (Q) secondary metabolites biosynthesis, transport, and catabolism; (T) signal transduction mechanisms; (U) intracellular trafficking, secretion, and vesicular transport; (V) defense mechanisms; (S) function unknown; (*) genes with no COG category assigned; following the respective colors in the legend. The outermost ring illustrates the G+C content.

### Core, accessory and unique genomes

The gene repertoire of CCBH4851 was compared to other strains divided in three groups (Table 1): (A) reference strains, (B) ST-277 strains, and (C) MDR strains belonging to other STs. A search for orthologs among these genomes identified genes that were shared by all genomes (core), genes specific to each genome (unique) or genes shared by two or more (but not all) genomes (accessory). In group A, 5,068 orthologous genes were identified as core genome, 251 as accessory genome and 703 genes were unique of CCBH4851 (S1 Table). Apart from genes included in the “Function unknown” COG class, the majority of unique genes were distributed among “Replication, recombination, and repair” and “Transcription” COG classes (Fig 2A). In group B, the core genome comprised 5,645 genes, which represents 80.52% of this group’s pangenome. Less than 20% were assigned to accessory and unique genomes, also having the “Replication, recombination, and repair” and “Transcription” COG classes as the major functional categories with exception of “Function unknown” family (Fig 2B). Group C, including MDR strains of different STs, presented 3,680 genes as core genome, 1,745 as accessory genome and 508 as unique genes of CCBH4851. The most evident functional categories remained the same (Fig 2C). The comparative analysis also revealed CCBH4851 genome had 205 genes lacking or presenting partial homology when compared to PAO1 genome (S2 Table). The predominant functional COG categories of the non-orthologous genes were “Replication, recombination, and repair”, “Transcription”, “Intracellular trafficking, secretion, and vesicular transport”, “Inorganic ion transport and metabolism”, “Energy production and conversion”, “Secondary metabolites biosynthesis, transport, and catabolism”, “Defense mechanisms”, and “Coenzyme transport and metabolism”. It is noteworthy that *oprD* (porin), *mexZ, liuR*, PA0207, PA0306, PA0306a, PA2100, PA2220, PA2547, PA3067, PA3094, and PA3508 (transcriptional regulators), *opdE* (membrane protein), *pilA* (type IV fimbrial protein), *algP* (alginate regulatory protein), and others were among these genes.

**Fig 2.**
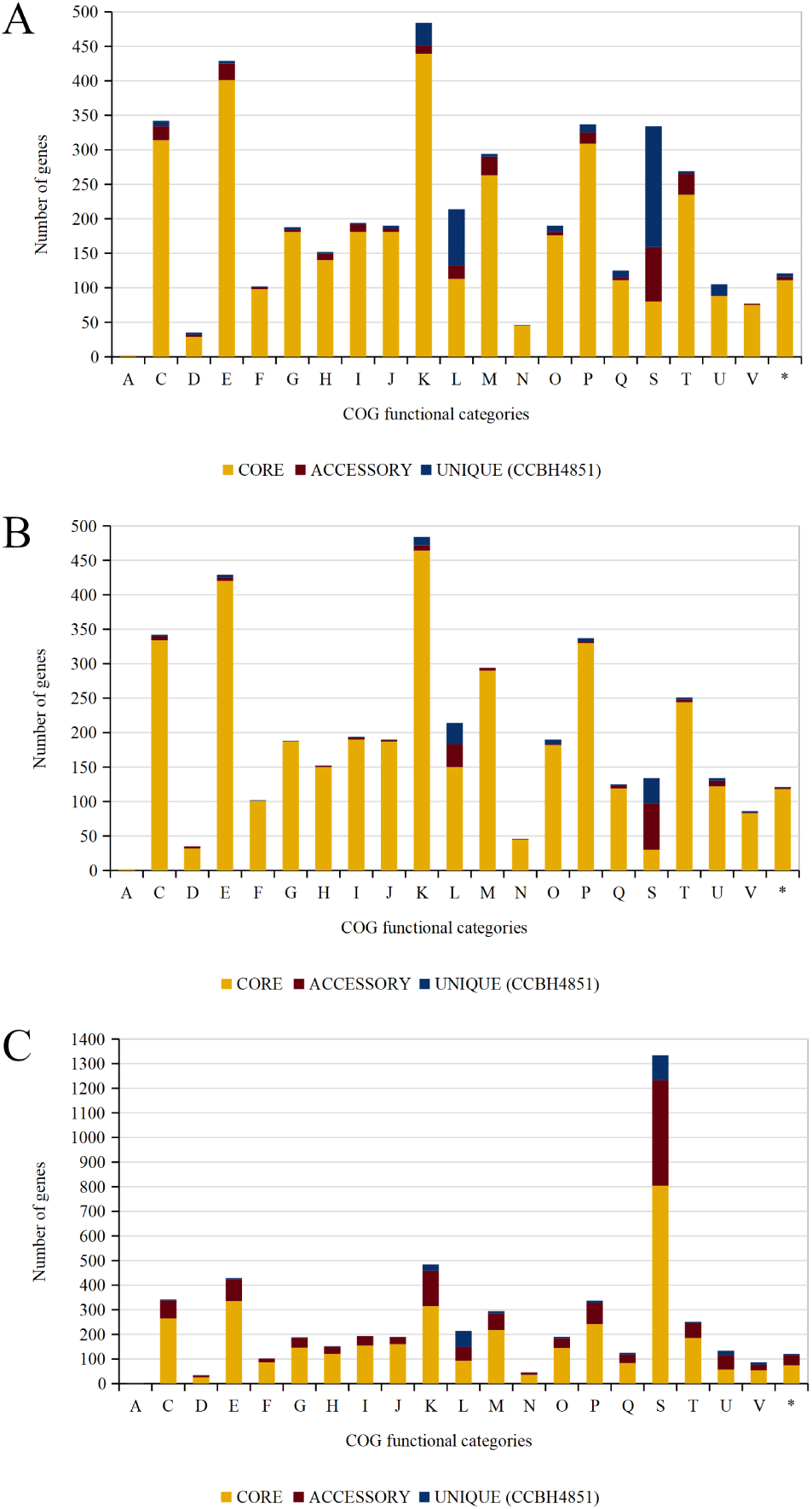
Orthology analysis and functional classification by COG. A: Core and accessory genomes shared between reference strains and CCBH4851, and unique genome of CCBH4851. B: Core and accessory genomes shared between other ST-277 strains and CCBH4851, and unique genome of CCBH4851. C: Core and accessory genomes shared between other ST strains and CCBH4851, and unique genome of CCBH4851. Legend of x-axis: (A) RNA processing and modification; (C) energy production and conversion; (D) cell cycle control, cell division, chromosome partitioning; (E) amino acid transport and metabolism; (F) nucleotide transport and metabolism; (G) carbohydrate transport and metabolism; (H) coenzyme transport and metabolism; (I) lipid transport and metabolism; (J) translation, ribosomal structure, and biogenesis; (K) transcription; (L) replication, recombination, and repair; (M) cell wall, membrane, and envelope biogenesis; (N) cell motility; (O) post-translational modification, protein turnover, and chaperones; (P) inorganic ion transport and metabolism; (Q) secondary metabolites biosynthesis, transport, and catabolism; (T) signal transduction mechanisms; (U) intracellular trafficking, secretion, and vesicular transport; (V) defense mechanisms; (S) function unknown; (*) genes with two or more COG categories assigned.

### Genomic islands and mobile elements

The *P. aeruginosa* CCBH4851 genome contained 19 integrated genomic islands with a total of 705 CDSs (Fig 2, yellow highlighted regions; S3 Table). Eighteen islands were homologous to *P. aeruginosa* genome islands (PAGIs) previously found in other clones belonging to ST-277 [25, 26]. An additional region found by IslandViewer was less than 5,000 bp, but had a G+C content of 55.13% and encoded 5 proteins including an integrase and an HTH domain-containing protein. This region was annotated as a new PAGI, numbered PAGI-41. Overall, 69.92% of the genes located in the PAGIs had no function assigned. The remainder were manly classified in the following COG categories: “Replication, recombination, and repair”, “Transcription”, “Intracellular trafficking, secretion, and vesicular transport”, “Energy production and conversion”, “Secondary metabolites biosynthesis, transport, and catabolism”, “Inorganic ion transport and metabolism” and “Defense mechanisms”. A total of 13 IS families were found in the CCBH4851 genome with 46 predicted interspersed open read frames (ORFs), including complete and partial sequences (S4 Table). Results of CRISPRCasFinder analysis revealed the presence of two CRISPR loci: one had only 1 spacer and direct repeats of 25 bp, and the other had 39 spacers and direct repeats of 32 bp. The first CRISPR was located at 1878269 to 1878370 genome position between two hypothetical protein-encoding genes (PA4851_08610 and PA4851_08615). The DNA sequence referring to this first CRISPR array was a variant of the intergenic region between PA3230 and PA3231 genes in PAO1 (homologous to PA4851_08610 and PA4851_08615). The second CRISPR was located at 5576114 to 5578733 genome position overlapping partially the hypothetical protein-encoding gene PA4851_25880. In addition, the second CRISPR array was located adjacent to an intact CRISPR-associated gene cluster, *cas2* (PA4851_25885), *cas1* (PA4851_25890), *cas4* (PA4851_25895), *cas7* (PA4851_25900), *cas8c* (PA4851_25905), *cas5* (PA4851_25910), *cas3* (PA4851_25915). These *cas* genes were related to the type I-C CRISPR-Cas system and were located inside the PAGI-34, as previously described in other ST-277 clones [25].

### Regulatory proteins

Comparative genome analysis revealed strong evidences of horizontal gene transfer events in CCBH4851 genome. Despite the predominance of genes encoding hypothetical proteins in these acquired regions, several CDSs were classified into the “Transcription” COG functional class. Regulatory protein analysis was performed to predict the presence of two-component systems, transcription factors, and other DNA-binding proteins in CCBH4851 genome. Table 3 summarizes a comparison between CCBH4851 and PAO1 regulatory proteins. In accordance with the BBH analysis, CCBH4851 genome was lacking the histidine-kinase PA2583, the response regulator PA0034, the transcriptional regulators PA3508, PA3067, and LiuR (PA2016). However, CCBH4851 genome possessed 28 additional regulatory proteins, mostly transcriptional regulators with Xre- and LysR-type domains, distributed along the PAGIs (S5 Table).

**Table 3.** Regulatory proteins of *P. aeruginosa* CCBH4851 compared to *P. aeruginosa* PAO1 reference strain.

| Strain | Predicted regulatory proteins |  |  |  |  |  |  |  |
| --- | --- | --- | --- | --- | --- | --- | --- | --- |
|  | TCS |  |  | TF |  |  |  | ODP |
|  | HK | RR | PP | TR | OCS | RR | SF |  |
| PAO1 | 62 | 73 | 4<sup>a</sup> | 170 | 185 | 49 | 24 | 36 |
| CCBH4851 | 61 | 72 | 5 | 189 | 191 | 48 | 25 | 38 |

TCS, two-component systems; TF, trascription factors; ODP, other DNA-binding proteins; HK, histidine-kinases; RR, response regulators; PP, phosphotransferase proteins; TR, transcriptional regulators; OCS, one-component systems; SF, sigma factors.
<sup>a</sup> A PP homologous sequence is present in the PAO1 DNA sequence, but it is not annotated in the deposited Genbank file.

### Antimicrobial resistance factors

Protein sequences of CCBH4851 genome were used for searches against CARD. As previously described [5], CCBH4851 genome possessed additional genes conferring multidrug resistance: two copies of *bla*_SPM-1_ (PA4851_13890, PA4851_13940), two copies of *sul1* (PA4851_12400, PA4851_12440), *rmtD* (PA4851_12415), *bla*_OXA-56_ (PA4851_12385), *aac(6’)-Ib7* (PA4851_12380), *aadA7* (PA4851_12390), *cmx* (PA4851_12450), and three copies of *bcr1* (PA4851_13870, PA4851_13920, PA4851_13970). As expected, all these genes were located into the PAGIs. Apart from *mexZ*, genes listed as PAO1 resistome were also present in CCBH4851 genome (Table 4).

**Table 4.** Resistance-associated genes of *P. aeruginosa* CCBH4851.

| Gene | Product | Location | Resistance mechanism |
| --- | --- | --- | --- |
| *ampC* | beta-lactamase AmpC | core genome | antibiotic inactivation |
| *aph* | aminoglycoside<br>3-N-acetyltransferase | core genome | antibiotic inactivation |
| *armR* | MexR antirepressor ArmR | core genome | antibiotic efflux |
| *arnA* | bifunctional UDP-glucuronic acid<br>decarboxylase/UDP-4-amino-4-deoxy-L-arabinose<br>formyltransferase | core genome | antibiotic target alteration |
| *arnC* | undecaprenyl-phosphate<br>4-deoxy-4-formamido-L-arabinose<br>transferase | core genome | antibiotic target alteration |
| *bacA* | undecaprenyl-diphosphatase | core genome | antibiotic target alteration |
| *cat* | chloramphenicol acetyltransferase | core genome | antibiotic inactivation |
| *cueR* | protein CueR | core genome | antibiotic efflux |
| *folA* | dihydrofolate reductase | core genome | antibiotic target replacement |
| *fusA1* | elongation factor G | core genome | antibiotic target alteration |
| *fusA2* | elongation factor G | core genome | antibiotic target alteration |
| *gyrA* | DNA gyrase subunit A | core genome | antibiotic target alteration |
| *kdpE* | two-component response regulator KdpE | core genome | antibiotic efflux |
| *mexA* | multidrug resistance protein MexA | core genome | antibiotic efflux |
| *mexB* | multidrug resistance protein MexB | core genome | antibiotic efflux |
| *mexC* | resistance-nodulation-cell division (RND) multidrug efflux membrane fusion protein MexC | core genome | antibiotic efflux |
| *mexD* | resistance-nodulation-cell division (RND) multidrug efflux transporter MexD | core genome | antibiotic efflux |
| *mexE* | resistance-nodulation-cell division (RND) multidrug efflux membrane fusion protein MexE | core genome | antibiotic efflux |
| *mexF* | resistance-nodulation-cell division (RND) multidrug efflux transporter MexF | core genome | antibiotic efflux |
| *mexG* | hypothetical protein | core genome | antibiotic efflux |
| *mexH* | resistance-nodulation-cell division (RND) efflux membrane fusion protein | core genome | antibiotic efflux |
| *mexI* | resistance-nodulation-cell division (RND) efflux transporter | core genome | antibiotic efflux |
| *mexR* | multidrug resistance operon repressor MexR | core genome | antibiotic efflux; antibiotic target alteration |
| *mexT* | transcriptional regulator MexT | core genome | antibiotic efflux |
| *nalC* | transcriptional regulator | core genome | antibiotic efflux |
| *nalD* | transcriptional regulator | core genome | antibiotic efflux |
| *nfxB* | transcriptional regulator NfxB | core genome | antibiotic efflux |
| *opmD* | hypothetical protein | core genome | antibiotic efflux |
| *oprJ* | multidrug efflux outer membrane protein OprJ | core genome | antibiotic efflux |
| *oprM* | outer membrane protein OprM | core genome | antibiotic efflux |
| *oprN* | multidrug efflux outer membrane protein OprN | core genome | antibiotic efflux |
| PA4851_00805 | resistance-nodulation-cell division (RND) efflux membrane fusion protein | core genome | antibiotic efflux |
| PA4851_00810 | resistance-nodulation-cell division (RND) efflux membrane fusion protein | core genome | antibiotic efflux |
| PA4851_00815 | resistance-nodulation-cell division (RND) efflux transporter | core genome | antibiotic efflux |
| PA4851_04160 | major facilitator superfamily transporter | core genome | antibiotic efflux |
| PA4851_06440 | transcriptional regulator | core genome | antibiotic efflux |
| PA4851_06445 | resistance-nodulation-cell division (RND) efflux membrane fusion protein | core genome | antibiotic efflux |
| PA4851_06450 | resistance-nodulation-cell division (RND) efflux transporter | core genome | antibiotic efflux |
| PA4851_07220 | resistance-nodulation-cell division (RND) efflux membrane fusion protein | core genome | antibiotic efflux |
| PA4851_07225 | resistance-nodulation-cell division (RND) efflux transporter | core genome | antibiotic efflux |
| PA4851_07230 | hypothetical protein | core genome | antibiotic efflux |
| PA4851_08745 | two-component response regulator | core genome | antibiotic efflux |
| PA4851_12380 | AAC(6')-Ib family aminoglycoside 6'-N-acetyltransferase | PAGI-25 | antibiotic inactivation |
| PA4851_12385 | OXA-10 family oxacillin-hydrolyzing class D beta-lactamase OXA-56 | PAGI-25 | antibiotic inactivation |
| PA4851_12390 | ANT(3'')-Ia family aminoglycoside nucleotidyltransferase AadA7 | PAGI-25 | antibiotic inactivation |
| PA4851_12415 | 16S rRNA (guanine(1405)-N(7))-methyltransferase RmtD1 | PAGI-25 | antibiotic target alteration |
| PA4851_12450 | chloramphenicol efflux MFS transporter Cmx | PAGI-25 | antibiotic efflux |
| PA4851_13685 | resistance-nodulation-cell division (RND) efflux transporter | core genome | antibiotic efflux |
| PA4851_13690 | resistance-nodulation-cell division (RND) efflux transporter | core genome | antibiotic efflux |
| PA4851_13695 | resistance-nodulation-cell division (RND) efflux transporter | core genome | antibiotic efflux |
| PA4851_13700 | hypothetical protein | core genome | antibiotic efflux |
| PA4851_13870 | putative chloramphenicol efflux MFS transporter Bcr1 | PAGI-15 | antibiotic efflux |
| PA4851_13920 | putative chloramphenicol efflux MFS transporter Bcr1 | PAGI-15 | antibiotic efflux |
| PA4851_13970 | putative chloramphenicol efflux MFS transporter Bcr1 | PAGI-15 | antibiotic efflux |
| PA4851_16855 | multidrug efflux lipoprotein | core genome | antibiotic efflux |
| PA4851_16860 | multidrug efflux protein | core genome | antibiotic efflux |
| PA4851_19800 | resistance-nodulation-cell division (RND) efflux transporter | core genome | antibiotic efflux |
| PA4851_19805 | resistance-nodulation-cell division (RND) efflux membrane fusion protein | core genome | antibiotic efflux |
| PA4851_20490 | multidrug resistance protein PmpM | core genome | antibiotic efflux |
| PA4851_21655 | glutathione transferase FosA | core genome | antibiotic inactivation |
| PA4851_24750 | resistance-nodulation-cell division (RND) efflux membrane fusion protein | core genome | antibiotic efflux |
| PA4851_24755 | multidrug efflux protein | core genome | antibiotic efflux |
| PA4851_26485 | transcriptional regulator | core genome | antibiotic efflux |
| PA4851_28475 | hypothetical protein | core genome | antibiotic efflux |
| PA4851_28555 | SMR multidrug efflux transporter | core genome | antibiotic efflux |
| PA4851_29765 | 2-octaprenyl-3-methyl-6-methoxy-1,4-benzoquinol hydroxylase | core genome | antibiotic inactivation |
| PA4851_31265 | beta-lactamase | core genome | antibiotic inactivation |
| PA4851_31285 | potassium efflux transporter | core genome | antibiotic efflux |
| *pmrA* | two-component regulator system response regulator PmrA | core genome | antibiotic target alteration |
| *pmrB* | two-component regulator system signal sensor kinase PmrB | core genome | antibiotic target alteration |
| *rpoB* | DNA-directed RNA polymerase subunit beta | core genome | antibiotic target alteration; antibiotic target replacement |
| *soxR* | redox-sensitive transcriptional activator SoxR | core genome | antibiotic efflux; antibiotic target alteration |
| *spm-1* | subclass B1 metallo-beta-lactamase SPM-1 | PAGI-15 | antibiotic inactivation |
| *spm-1* | subclass B1 metallo-beta-lactamase SPM-1 | PAGI-15 | antibiotic inactivation |
| *str* | streptomycin 3"-phosphotransferase | core genome | antibiotic inactivation |
| *sul1* | sulfonamide-resistant dihydropteroate synthase Sul1 | PAGI-25 | antibiotic target replacement |
| *sul1* | sulfonamide-resistant dihydropteroate synthase Sul1 | PAGI-25 | antibiotic target replacement |
| *tufA* | elongation factor Tu | core genome | antibiotic target alteration |
| *tufB* | elongation factor Tu | core genome | antibiotic target alteration |
| *vfr* | cAMP-regulatory protein | core genome | antibiotic efflux |

### Single nucleotide polymorphisms

SNP calling between *P. aeruginosa* CCBH4851 and PAO1 genome identified 25,220 variant types in the genome including SNPs *per se*, multiple nucleotide polymorphisms (MNPs), insertions, deletions and a combination of SNP/MNP [23]. Results revealed 16,690 synonymous variants, 4,972 missense variants, 7 stop gained, 5 stop lost, 5 start lost, 39 in frame deletions/insertions, 40 frameshift variants, and 3440 variant types affecting intergenic regions (S6 Table). The amount of synonymous variants was left aside as they have no presumable impact on cellular processes. Apart from intergenic regions, the remainder affected genes were classified into COG families to assess whether variant types were common to a few functional categories or were randomly distributed among all of them. Mutations occurred mainly in genes belonging to “Inorganic ion transport, and metabolism”, “Amino acid transport and metabolism”, “Cell wall, membrane, and envelope biogenesis”, “Transcription”, and “Energy production and conversion” COG categories (Fig 3). In addition, Table 5 summarizes a list of virulence and antimicrobial resistance-associated genes with their respective amino acid substitutions. The structural effect of these substitutions was predicted and described in details in S7 Table.

**Table 5.**
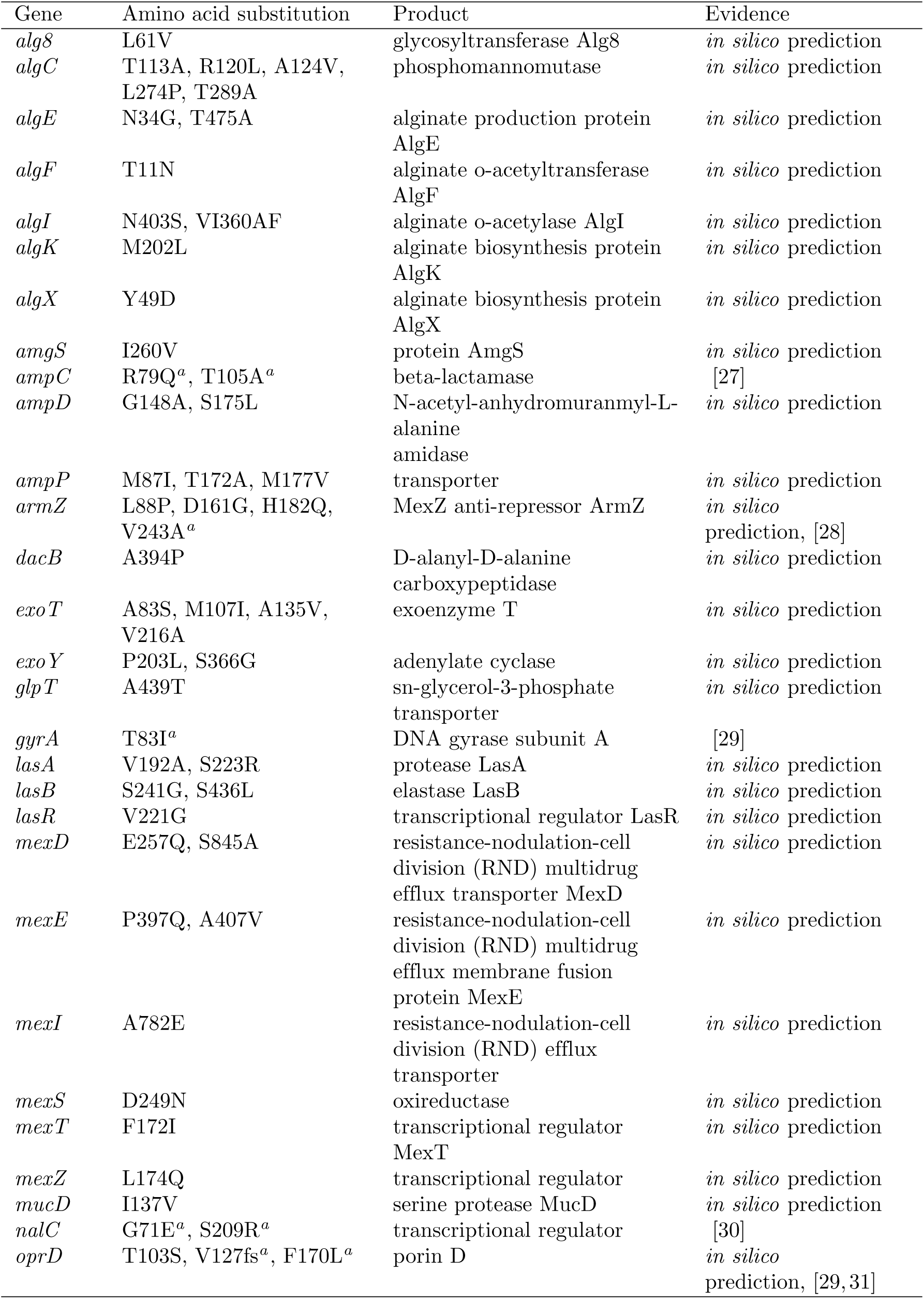
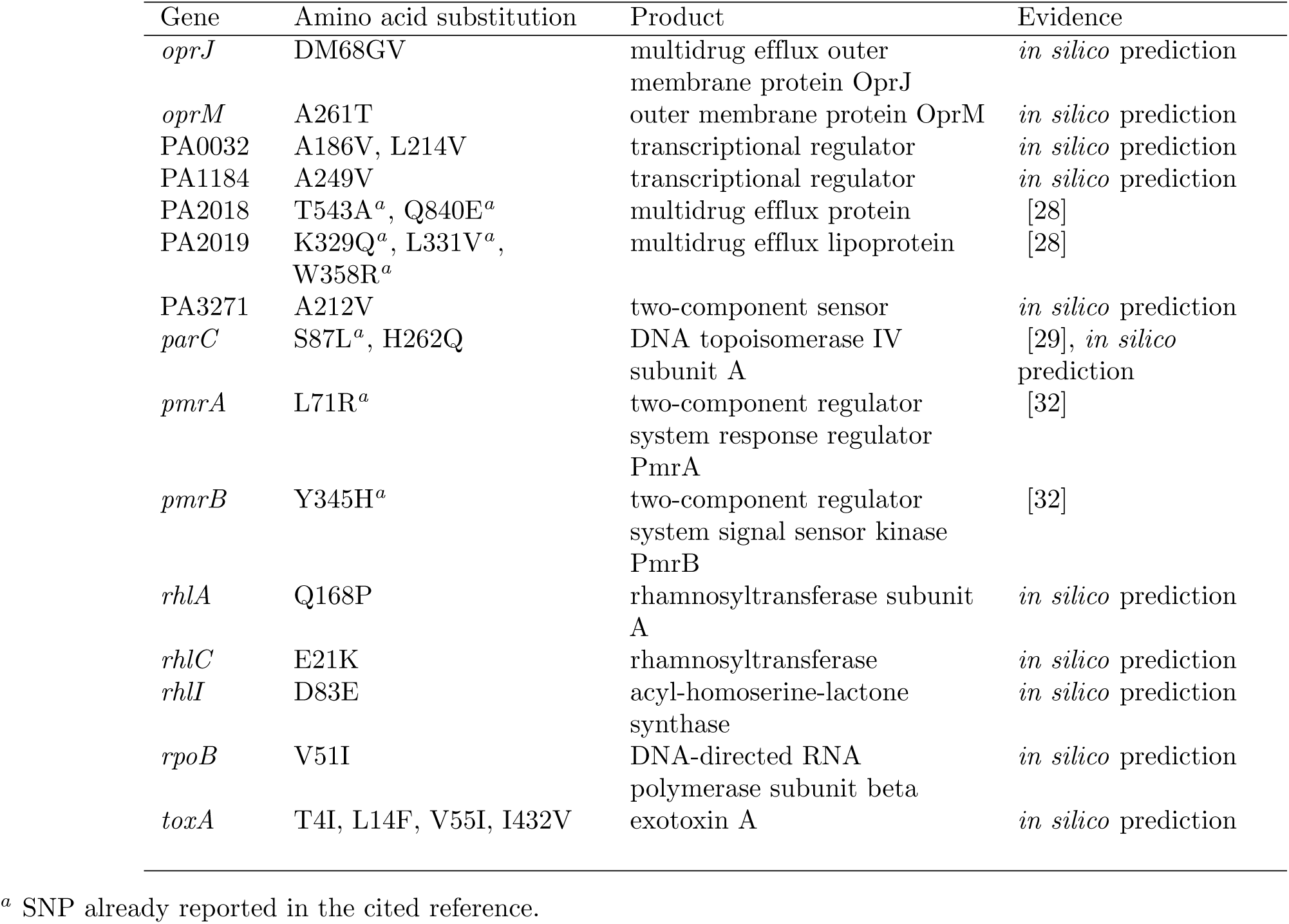
SNPs found in virulence and antimicrobial resistance-associated genes of *P. aeruginosa* CCBH4851 using as reference *P. aeruginosa* PAO1.

**Fig 3.**
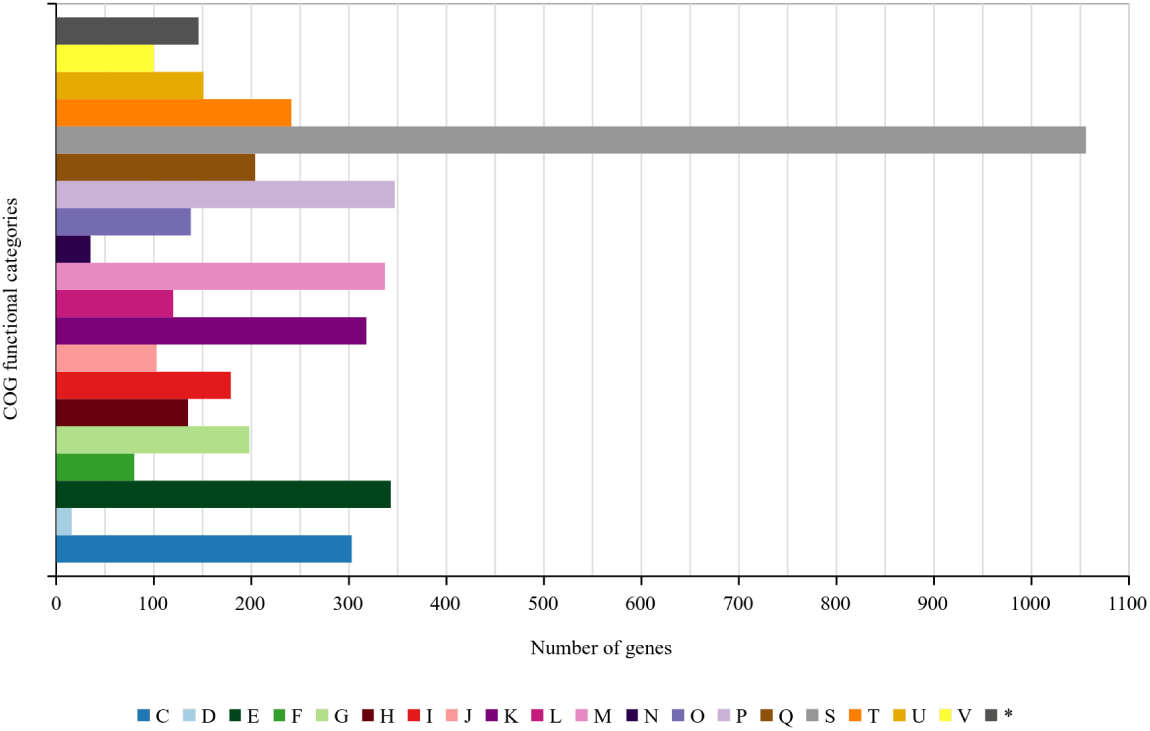
Functionally clustered genes affected by SNPs causing amino acid substitutions. Description of COG categories: (C) energy production and conversion; (D) cell cycle control, cell division, chromosome partitioning; (E) amino acid transport and metabolism; (F) nucleotide transport and metabolism; (G) carbohydrate transport and metabolism; (H) coenzyme transport and metabolism; (I) lipid transport and metabolism; (J) translation, ribosomal structure, and biogenesis; (K) transcription; (L) replication, recombination, and repair; (M) cell wall, membrane, and envelope biogenesis; (N) cell motility; (O) post-translational modification, protein turnover, and chaperones; (P) inorganic ion transport and metabolism; (Q) secondary metabolites biosynthesis, transport, and catabolism; (S) function unknown; (T) signal transduction mechanisms; (U) intracellular trafficking, secretion, and vesicular transport; (V) defense mechanisms; (*) genes with two or more COG categories assigned; following the respective colors in the legend.

## Discussion

*Pseudomonas aeruginosa* CCBH4851 is a clinical isolate originating from Brazil, which belongs to the endemic group ST-277. Previous studies demonstrate ST277 strains have a highly conserved DNA sequence and share several virulence and antimicrobial resistance-related features. However, to date, the presence of *bla*_SPM-1_ gene conferring resistance to carbapenem is a singular trait of Brazilian clones [4, 25]. The purpose of this study was to reassemble and to re-annotate CCBH4851 genome in order to perform a thorough characterization and a genomic comparison with other *P. aeruginosa* strains. The reassembly resulted in a single contig representing the complete closed genome of CCBH4851. Together with the provided annotation, data such as chromosome size, G+C content, number of CDSs, structural RNAs, and tRNAs, were consistent with general features reported for *P. aeruginosa* strains [33]. In addition, CCBH4851 genome alignment with PAO1, PA14, and ST-277 strains revealed a strong synteny (S1 Fig) indicating an accurate assembly and annotation. Corroborating this synteny, the core genome of CCBH4851, PAO1 and PA14 strains covered more than 80% of CCBH4851 genome. A similar percentage was found when the core genome was defined comparing CCBH4851 with other ST-277 clones. However, when comparing MDR strains belonging to other STs, the core genome was smaller and an increase in the number of genes comprised in the accessory genome was observed. This result could be attributed to different factors: (i) some strains used in this analysis had more than hundreds contigs and this lack of continuity could affect genome annotation; (ii) the strains used in this analysis are from different countries and had distinct resistance profiles, which could suggest a variation of mutation patterns and acquired regions, depicting interstrain differences. The larger accessory genome among these strains suggests a variability in pathogenicity and environmental flexibility. The comparative genome analysis revealed total or partial absence of homology between several PAO1 genes and CCBH4851 genome. Among them, there is *oprD*, which suffered a deletion of 2 nucleotides in the 380-381 positions causing a frameshift resulting in the gain of a premature stop codon. OprD is an outer-membrane porin important for the diffusion of carbapenems, particularly imipenem. Disruption of OprD is a common resistance mechanism, manly when combined with overexpression of AmpC and efflux pump systems [31]. Indeed, the mutation of *mexZ* observed in CCBH4851 suggests the overexpression of at least one efflux pump system, MexXY, whose expression is repressed by MexZ. A deletion of 17 nucleotides from the position 439 caused a frameshift resulting in the stop codon loss. Mutations in *mexZ* are often present in clinical isolates overexpressing MexXY [30]. Other differences revealed by the comparative analysis were the mutations of *algP* and *pilA* genes, both related to biofilm development. AlgP is a histone-like protein which activates the transcription of *algD*, responsible for the alginate precursor synthesis. The amino acid sequence of AlgP contains several repeated KPAA motifs which are mutation targets at high frequency in clinical isolates. According to this, the *algP* gene sequence of CCBH4851 had an insertion of 12 nucleotides, adding one KPAA motif to the AlgP protein sequence. The repeated 12-bp sequences of *algP* are considered a hot spot for DNA rearrangements and could provide a reversible switching mechanism between nonmucoid and mucoid phenotypes, thus turning on and off virulence factors important for a successful infection [34]. On the other hand, *pilA* sequence is completely absent from CCBH4851 genome. PilA is a major pilin protein involved in type IV pilus (T4P) biogenesis. Studies demonstrate PilA loss contributes to a decrease in cyclic AMP intracellular levels and to an increased expression of a set of genes, such as *hcpA* and *hcpB*, which are involved in the type VI secretion system (T6SS). T6SS are related to virulence, biofilm formation and biofilm-specific antibiotic resistance. However, PilA mutants are T4P defective, which could hinder biofilm formation. *In vitro* analysis showed biofilm formation occurs differently under distinct conditions and biofilm development is possible independent of the flagella or T4P presence. Although the impact of PilA loss is not fully understood, the absence of *pilA* in carbapenem-resistant clinical isolates is not uncommon [35–37]. The acquisition of exogenous material by horizontal gene transfer is a common adaptive mechanism among *P. aeruginosa* strains, often related to the presence pf genomic islands. Genomic islands are clusters of genes often encoding virulence factors, antimicrobial resistance proteins, toxins, secretion system proteins, transcriptional regulators, and other proteins [38]. CCBH4851 genome presented a high number of islands, all sharing homology with previously described PAGIs, except for the PAGI-41. The majority of proteins encoded by these PAGIs are hypothetical proteins, however, some of them could be clustered in functional categories due the presence of conserved domains. The outstanding categories comprised genes involved in mechanisms of replication, recombination and repair, and transcription. Indeed, the newly annotated PAGI-41 carries an XRE-type HTH domain-containing protein, which is usually a repressor involved in the metabolism of xenobiotics. Moreover, all additional genes conferring resistance to a broad range of antimicrobial agents were located in the PAGIs, including the two copies of the carbapenem resistance gene *bla*_SPM-1_. In addition to the presence of PAGIs, distinct IS families were also detected in the CCBH4851 genome. The importance of ISs is not restricted to their role in the horizontal gene transfer, but ISs movement along the chromosome can also affect antibiotic resistance by the activation of gene expression [39]. Another player in shaping the bacterial genome is the CRISPR-Cas system. The type I-C found in CCBH4851 genome was previously described in other ST-277 bacteria as well as in members of ST-235 [25, 40]. A recent study suggests the CRISPR-Cas systems’ protective effect is more evident at the population level than at an evolutionary scale [41]. This could explain the intraclonal genome conservation observed among Brazilian isolates belonging to ST-277 [25]. During the infection course, *P. aeruginosa* strains tend to adapt to the selective pressures of the environment, often through mutations in intergenic and/or coding sequence regions. A classical example of adaptive mutation is the MexT protein, responsible for activation of the *mexEF-OprN* operon. MexEF-OprN is quiescent in wild-type cells and its expression occurs following mutations in MexT or MexT-related genes. Indeed, the *mexT* gene sequence of CCBH4851 had an 8-bp deletion known to render MexT active, causing the induction of *mexEF-oprN* transcription and a decrease in OprD levels, which characterizes the so-called *nfxC* -type mutants [42]. However, later work showed that only this 8-bp deletion was not enough to activate the transcription of *mexE* and the generation of *nfxC* -type mutants seems to be multifactorial. Indeed, *mexS* inactivation, a gene upstream of *mexT*, seems to be one of these additional factors [43]. The *mexS* gene of CCBH4851 suffered a deletion of 1 nucleotide in position 22 causing a frameshift which resulted in the stop codon loss. The gene product of the *mexS* pseudogene showed an alignment of only 7 aa with the MexS wild-type sequence of PAO1. This could suggest the absence of MexS in CCBH4851 proteome, leading to the overexpression of *mexE* and other phenotypes related to *nfxC* -type mutants. Table 5 summarizes other SNPs in virulence and resistance-associated genes that are frequently mutated in *P. aeruginosa* clinical isolates. Among them, amino acid substitutions T83I in *gyrA* and S87L in *parC* are well-known to play an important role in fluoroquinolone resistance as well as the overexpression of efflux pump systems. Although quantification of efflux pump systems expression is required to confirm altered transcription levels, amino acid substitutions in *nalC* (repressor of *mexA* gene), *armZ* (MexZ anti-repressor), PA2018 and PA2019 suggest additional mechanisms leading to the overexpression of *mexA* and *mexX*. In addition to these factors, SNPs in the *ampC* gene of CCBH4751 cause a modification in AmpC protein which characterizes it as the variant type PDC-5. Clinical isolates carrying this variant presented AmpC overexpression, increased beta-lactamase activity and reduced susceptibility to ceftazidime, cefepime, cefpirome, aztreonam, imipenem, and meropenem [27–30]. Not all SNPs found in this work were previously described; however, different mutations in the same genes listed in Table 5 were observed in other clinical isolates, suggesting a recurrence in the mutations’ location caused by selective pressure. The variants found in CCBH4851 could be intraclonal-specific, but could still change gene function, thus contributing to the enhanced resistance of ST-277 clones. In fact, *in silico* analysis suggest some of these SNPs could affect the protein function due the introduction of amino acids with different properties, such as size, charge, hydrophobicity, often located in protein domains. The prediction indicates noteworthy mutations occurring in genes such as *ampD* (negative regulator of *ampC* expression), *mucD* (repressor of *algU* alternative sigma factor transcription), *oprJ* (member of *mexCD* efflux system), and others; which could affect the protein folding and/or function (S7 Table).

## Conclusion

The features found in the *P. aeruginosa* CCBH4851 complete genome are somehow related to its pathoadaptive behavior, since part of them are common among clinical isolates and some are unique of ST-277 clones. The number of unique genes, the number of genes contained in the acquired genomic islands, and the larger size of CCBH4851 chromosome are consistent with this observation. However, the majority of the genome is shared with more susceptible strains, suggesting the high number of mutations found in conserved genes are contributing to the success of this MDR strain. Further validation of uncharacterized polymorphisms revealed by this study should help to increase the understanding of CCBH4851 phenotypes. The genome characterization and the comparative analysis presented here provide some insights into bacterial virulence and antibiotic resistance mechanisms that may contribute to the future development of therapeutic choices in *P. aeruginosa* infections.

## Supporting information

**S1 Fig. Pairwise alignment between *Pseudomonas aeruginosa* chromosomes.** Coloured blocks indicate conserved and highly related genomic regions. Blocks shifted below the horizontal axis indicate segments that align in the reverse orientation relative to the reference strain *Pseudomonas aeruginosa* PAO1.

**S1 Table. Unique genes of *Pseudomonas aeruginosa* CCBH4851 compared to reference strains *Pseudomonas aeruginosa* PAO1 and PA14.**

**S2 Table. List of *Pseudomonas aeruginosa* PAO1 genes (with assigned function) presenting partial or no homology with *Pseudomonas aeruginosa* CCBH4851 genome.**

**S3 Table. Genomic islands found in the *Pseudomonas aeruginosa* CCBH4851 closed genome.**

**S4 Table. Insertion sequence families and predicted ORFs found in the *Pseudomonas aeruginosa* CCBH4851 complete genome.**

**S5 Table. Predicted regulatory proteins of *Pseudomonas aeruginosa* CCBH4851.**

**S6 Table. Single nucleotide polymorphism variants found in the *Pseudomonas aeruginosa* CCBH4851 using as reference *Pseudomonas aeruginosa* PAO1.**

**S7 Table. Structural effect of amino acid substitutions in virulence and antimicrobial resistance-associated genes of *Pseudomonas aeruginosa* CCBH4851 predicted by HOPE web application.**

## Acknowledgments

This work was supported by INOVA-FIOCRUZ (grant no. VPPCB-007-FIO-18-2-29) and FAPERJ (grant no. E-26/202.763/2015).

