## Supplementary figures and images for "Comparative genome analysis of a multidrug-resistant *Pseudomonas aeruginosa* sequence type 277 clone that harbours two copies of the *bla*_SPM-1_ gene and multiple single nucleotide polymorphisms in other resistance-associated genes"

### S1 Fig

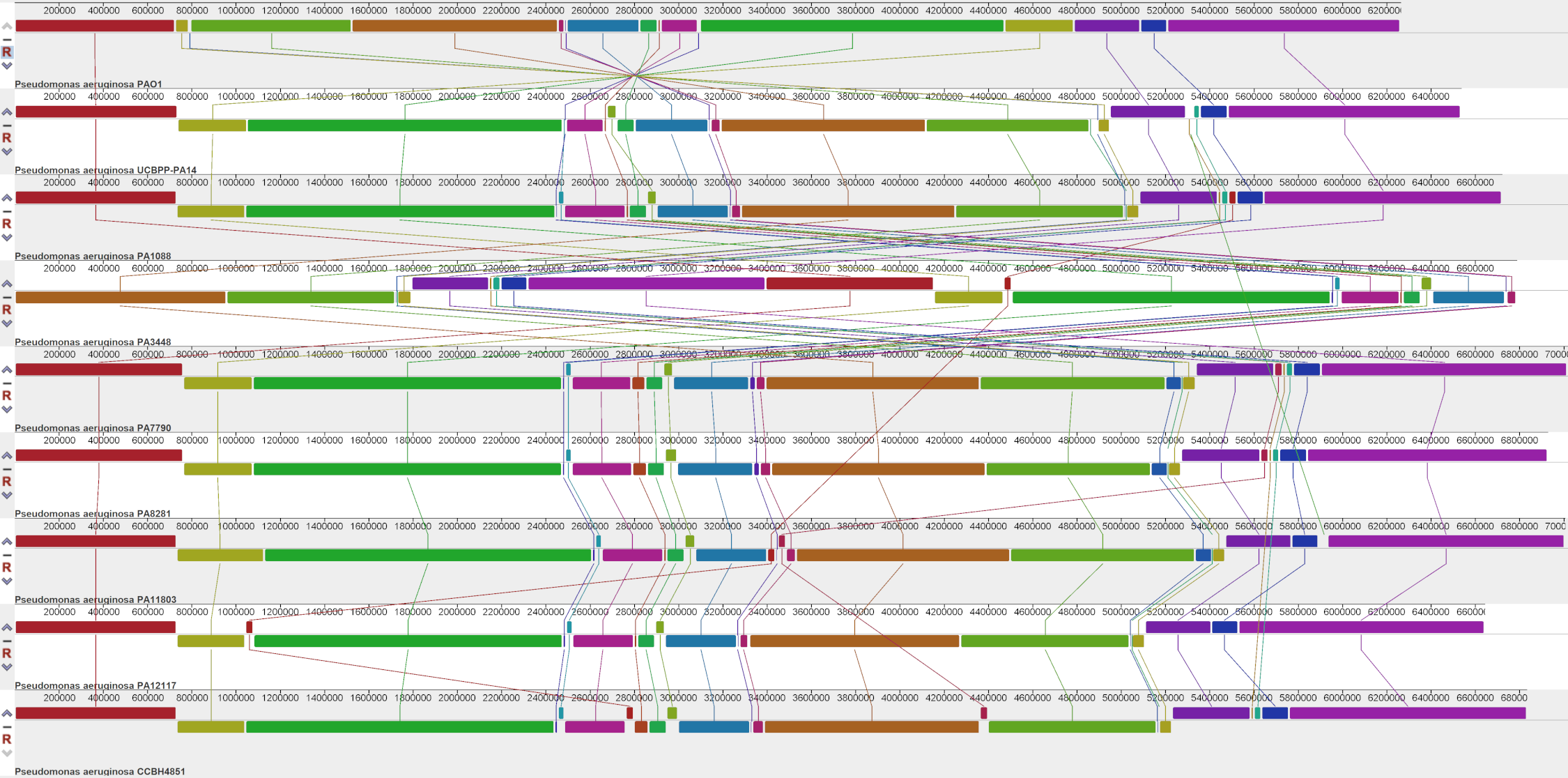
